# The eEF2 kinase coordinates the DNA damage response to cisplatin by supporting p53 activation

**DOI:** 10.1101/2023.03.28.534603

**Authors:** Jonathan KM Lim, Arash Samiei, Christopher J Carnie, Vanessa Brinkman, Daniel Radiloff, Jordan Cran, Gabriel Leprivier, Poul H Sorensen

**Affiliations:** Institute of Neuropathology, University Hospital Düsseldorf, Medical Faculty, Heinrich Heine University, Düsseldorf, Germany; Department of Pathology and Laboratory Medicine, University of British Columbia, Vancouver, British Columbia, Canada; Department of Molecular Oncology, BC Cancer Research Institute, Vancouver, British Columbia, Canada; The Wellcome Trust and Cancer Research UK Gurdon Institute, and Department of Biochemistry, University of Cambridge, Cambridge, UK; Institute of Toxicology, Heinrich Heine University, Düsseldorf, Germany; Terry Fox Laboratory, BC Cancer Research Institute, Vancouver, British Columbia, Canada

## Abstract

Eukaryotic elongation factor 2 (eEF2) kinase (eEF2K) is a stress-responsive hub that inhibits the translation elongation factor eEF2, and consequently mRNA translation elongation, in response to hypoxia and nutrient deprivation. EEF2K is also involved in the response to DNA damage but its role in response to DNA crosslinks, as induced by cisplatin, is not known. Here we found that eEF2K is critical to mediate the cellular response to cisplatin. We uncovered that eEF2K deficient cells are more resistant to cisplatin treatment. Mechanistically, eEF2K deficiency blunts the activation of the DNA damage response associated ATM and ATR pathways, in turn preventing p53 activation and therefore compromising induction of cisplatin-induced apoptosis. We also report that loss of eEF2K delays the resolution of DNA damage triggered by cisplatin, suggesting that eEF2K contributes to DNA damage repair in response to cisplatin. In support of this, our data shows that eEF2K promotes the expression of the DNA repair protein ERCC1, critical for the repair of cisplatin-caused DNA damage. Finally, using *Caenorhabditis elegans* as an in vivo model, we find that deletion of *efk-1*, the worm *eEF2K* ortholog, mitigates the induction of germ cell death in response to cisplatin. Together, our data highlight that eEF2K represents an evolutionary conserved mediator of the DNA damage response to cisplatin which promotes p53 activation to induce cell death, or alternatively facilitates DNA repair, depending on the extent of DNA damage.

## INTRODUCTION

Eukaryotic elongation factor 2 (eEF2) kinase (eEF2K) is an atypical protein kinase that acts as an evolutionarily conserved regulator of protein synthesis. It does so by inhibiting the elongation step of mRNA translation via phosphorylation and inactivation of its thus far exclusive substrate, eEF2 [1]. Previous studies illustrate a cytoprotective role of eEF2K, since it mediates the cellular response to various forms of stress including nutrient deprivation, hypoxia, and DNA damage. Notably, eEF2K is a downstream effector of AMP activated protein kinase (AMPK) and mammalian target of rapamycin complex 1 (mTORC1) pathways, both master regulators of the cellular response to metabolic and energetic stress [2]. Indeed, we previously showed that eEF2K protects mammalian cells as well as the nematode worm *Caenorhabditis elegans* (*C. elegans*) from nutrient depletion by blocking translation elongation, and that this pathway is exploited by cancer cells during their adaptation to such metabolic stress [3, 4]. Moreover, eEF2K promotes the survival of cancer cells and primary neurons under hypoxia through a similar mechanism [5, 6].

In addition, eEF2K is involved in the DNA damage response (DDR), exhibiting both protective and pro-apoptotic roles. Namely, it was reported that in response to doxorubicin treatment, AMPK-mediated activation of eEF2K induces stalling of translation elongation, whereas upon recovery from such genotoxic stress, eEF2K is specifically targeted for proteosomal degradation via its interaction with the Skp1–cullin–F-box protein (SCF)-beta-transducin repeats-containing proteins (βTrCP) ubiquitin ligase complex, allowing resumption of translation elongation [7]. In relation to this, it was shown that eEF2K maintains germline quality by inducing apoptosis of defective germ cells and oocytes following exposure to hydrogen peroxide and doxorubicin both in *C. elegans* and mice, with eEF2K loss resulting in accumulation of poor-quality oocytes and a decline in embryonic viability [8]. Similarly, eEF2K knockout in mice reduced apoptosis in bone marrow and gastrointestinal tracts following ionizing radiation [9]. Paradoxically, eEF2K deficiency led to more pronounced DNA damage and mitotic cell death in small intestinal stem cells and conversely, overexpression of eEF2K in cancer cells decreased radiation-induced apoptosis [9]. Thus, whether eEF2K mediates cellular and tissue protection or conversely, drives cellular susceptibility to genotoxic stress is still unclear. Moreover, how eEF2K responds to other inducers of DNA damage and the exact role of eEF2K in modulating the DDR is not well understood.

One typical DNA damage inducer is cisplatin, which belongs to the platinum group of chemotherapeutic drugs and also considered an alkylating-like agent. The cytotoxic effect of cisplatin arises through the formation of intra- and inter-strand DNA crosslinks (ICLs), and DNA-protein crosslinks [10]. The interaction of cisplatin with DNA blocks the progression of replicative DNA polymerases during S-phase, which may lead to the formation of double-strand breaks (DSBs) that are a highly toxic form of DNA damage due to their propensity to give rise to insertions, deletions or even translocations [11, 12]. The repair of ICLs is complex, involving multiple interacting pathways and protein mediators. Namely, the nucleotide excision repair (NER) pathway is the best understood mechanism employed by the cell to repair intra-strand crosslink adducts as well as ICLs [11]. While most NER proteins are associated with replication-independent ICL repair, the DNA excision repair endonuclease ERCC1-XPF complex also plays a prominent role in replication-dependent ICL repair by interacting with the Fanconi anemia (FA) and homologous replication repair (HRR) pathways, and is described to reduce DSB levels after cisplatin treatment [13, 14, 15]. Given the clinical relevance of eEF2K in promoting cancer cell survival and tumour development, and the use of cisplatin as a chemotherapeutic agent for multiple tumour types, we sought to clarify the relationship between the molecular functions of eEF2K and cisplatin-induced DNA damage. Notably, the role of eEF2K in the response to cisplatin is unknown.

Here, we show that eEF2K mediates the susceptibility of cells towards cisplatin, since eEF2K deficient cell lines exhibit reduced levels of apoptosis under such conditions. Moreover, we report that the worm eEF2K ortholog, *efk-1*, mediates germ cell death in *C. elegans* in response to cisplatin. Interestingly, while eEF2K supports the induction of the DDR pathway and promotes enhanced repair in response to cisplatin, it also leads to p53 activation and expression, thereby mediating DNA damage mediated cell death under chronic cisplatin treatment. Overall, our findings illustrate a critical role of eEF2K in mounting a DDR to cisplatin but also in driving cell death, presumably to safeguard against the propagation of deleterious effects from DNA damage.

## RESULTS

### eEF2K deficiency decreases susceptibility of cells to cisplatin

We first asked what roles eEF2K might play in the cellular response to DNA damage induced by cisplatin. To investigate this, we subjected *Eef2k*-knockout (*Eef2k*^-/-^) mouse embryonic fibroblasts (MEFs) and wild type control MEFs (*Eef2k*^+/+^; Figure S1A) to increasing concentrations of cisplatin treatment, and assessed cell viability using an MTT assay. While both cell lines displayed decreased viability in response to cisplatin, *Eef2k*^-/-^ cells exhibited reduced susceptibility to this agent as compared to *Eef2k*^+/+^ cells (Figure 1A). To recapitulate this finding in a different cell model, we generated HEK293 cells that express two distinct short hairpin RNAs (shRNAs) targeting eEF2K mRNA (sh-eEF2K1 and 2; Figure S1B), that were similarly exposed to cisplatin treatment followed by MTT assays. Notably, sh-eEF2K cells showed a minimal decrease in viability under these conditions compared to control cells (sh-scr), which in contrast were highly susceptible to cisplatin treatment (Figure 1B). Additionally, the sensitivity of HEK293 cells to cisplatin was accompanied with induction of eEF2 phosphorylation in response to cisplatin (Figure S1C). To determine if the decreased viability of eEF2K-competent cells (i.e. *Eef2k*^+/+^ and sh-scr HEK293 cells) treated with cisplatin was due to induction of cell death or a decrease in proliferation, we exposed *Eef2k*^+/+^ and *Eef2k*^-/-^ cells to cisplatin and quantified cell death using trypan blue-exclusion assays. Consistent with the viability assays, *Eef2k*^+/+^ cells displayed increased cell death relative to *Eef2k*^-/-^ cells over a broad concentration range (Figure 1C). Moreover, we ascertained that this increase in cell death was consistent with induction of apoptosis, since *Eef2k*^-/-^ cells showed markedly less cleaved-PARP and cleaved-Caspase-3 levels following cisplatin treatment, as compared to *Eef2k*^+/+^ cells (Figure 1D). This pattern was confirmed in sh-eEF2K versus sh-scr HEK293 cells treated with cisplatin (Figure 1E). Together, these data suggest that eEF2K mediates the susceptibility of cells towards cisplatin, and raise the possibility that eEF2K deficiency results in either more proficient repair of cisplatin-induced DNA damage, or dampened DNA damage-induced pro-apoptotic signalling.

**Figure 1.**
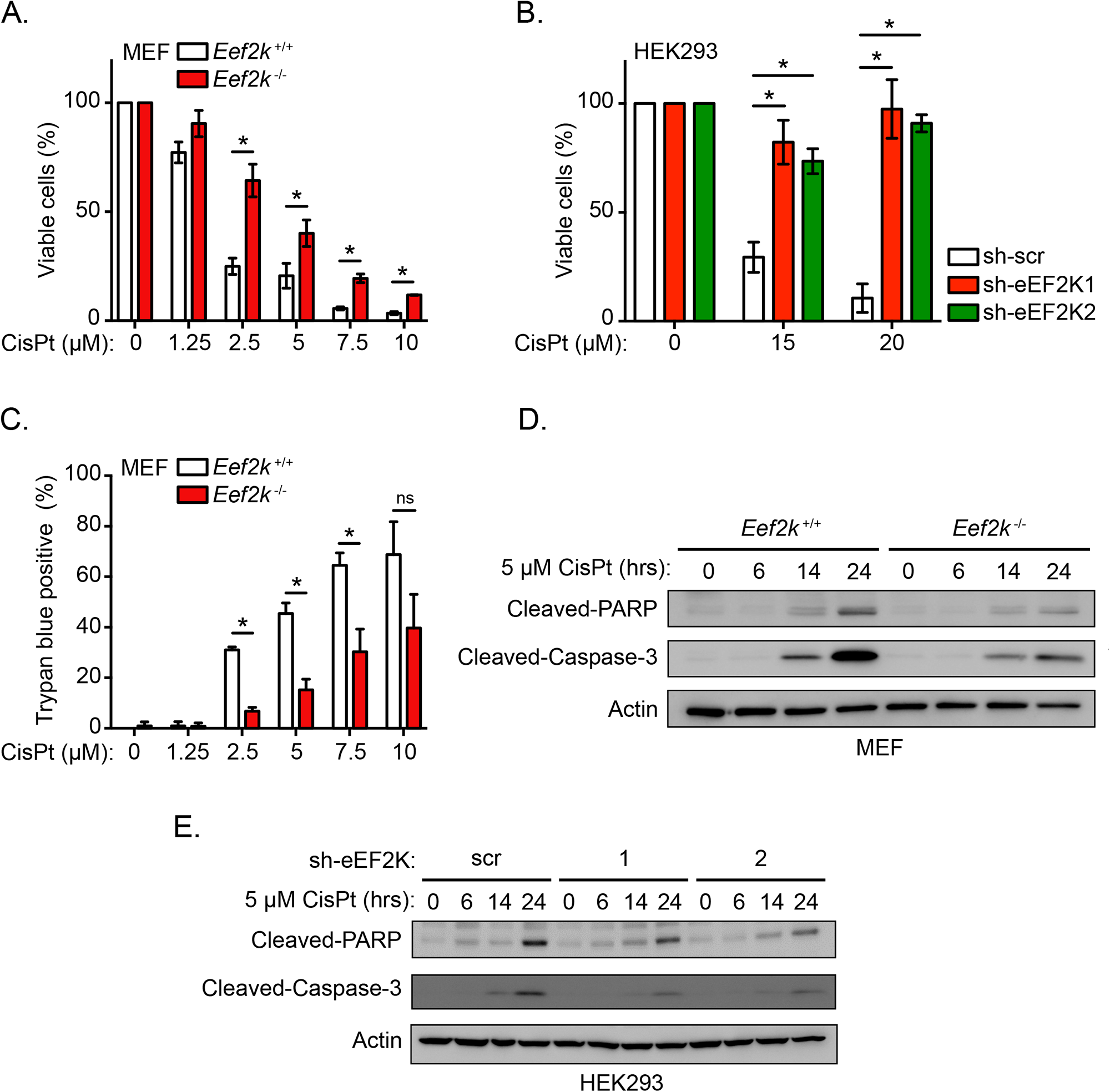
eEF2K mediates cellular susceptibility towards cisplatin. **A.** Viability of *Eef2k*^+/+^ and *Eef2k*^-/-^ MEFs treated with the indicated concentrations of cisplatin (CisPt) or vehicle control for 48 hours as measured by an MTT assay (n = 3). **B.** Viability of HEK293 cells stably expressing individual shRNAs targeting eEF2K (sh-eEF2K1 and sh-eEF2K2) or scrambled control (sh-scr) treated with the indicated concentrations of CisPt or vehicle control for 72 hours as measured by an MTT assay (n = 3). **C.** Cell death rates of *Eef2k*^+/+^ and *Eef2k*^-/-^ MEFs treated with the indicated concentrations of CisPt or vehicle control for 48 hours as indicated, as measured using Trypan blue staining (n = 3). **D.** Level of apoptosis markers in *Eef2k*^+/+^ and *Eef2k*^-/-^ MEFs treated with CisPt (5 μM) for the indicated times, as measured with immunoblot analysis of cleaved-PARP and cleaved-Caspase-3. **E.** Level of apoptosis markers in HEK293 cells stably expressing sh-eEF2K1, sh-eEF2K2, or sh-scr treated with CisPt (5 μM) for the indicated times, as measured with immunoblot analysis of cleaved-PARP and cleaved-Caspase-3. Data are expressed as mean ± SD; \**P* < 0.05, ns = non-significant.

### eEF2K contributes to the induction of DDR pathways in response to cisplatin

Cisplatin-induced cytotoxicity is primarily mediated by cisplatin-DNA adducts and ICLs, which leads to the formation of stalled replication forks, engagement of the Fanconi anemia pathway and generation of DNA double-stranded breaks (DSBs). Since these forms of DNA damage converge on ataxia-telangiectasia mutated (ATM) and ATM- and Rad3-Related (ATR) kinases (Figure 2A), we examined DDR signalling downstream of these kinases in cisplatin-treated *Eef2k*^+/+^ and *Eef2k*^-/-^ cells. We found that the Atm and Atr-mediated DDR was compromised in *Eef2k*^-/-^ cells, as indicated by reduced activation of Atm (phosphorylation at Ser1981) and Atr (phosphorylation at Ser428), reduced activation of Chk1 (phosphorylation at Ser345) and p53 (phosphorylation at Ser15), as well as reduced phosphorylation of H2AX (known as γH2AX) and p21 levels under cisplatin treatment over time (Figure 2B). Moreover, *Eef2k*^-/-^ cells exhibited reduced total levels of Atm, Atr, Chk1, and p53 compared to *Eef2k*^+/+^, suggesting that the blunted DDR in these cells is partially owed to reduced availability of these protein effectors. To eliminate the possibility that the compromised DDR pathways in *Eef2k*^-/-^ cells occurs as an adaptive process to genetic loss of eEF2K, we also used small interfering RNA (siRNA) to acutely target eEF2K in wild type MEFs under cisplatin-treatment. Similar to *Eef2k*^-/-^ cells, si-eEF2K cells displayed reduction of ATM and ATR activation, significant decreases in CHK1 and p53 activation, as well as a significant reduction in γH2AX and p21 levels relative to control cells (scr) under cisplatin treatment over a time course (Figure 2C). Notably, there were no observable differences in total ATM and ATR levels in these cells, and only marginal reduction in total CHK1 and p53 levels in si-eEF2K cells relative to control cells. The basis of this discrepancy compared to the knockout cells remains unknown.

**Figure 2.**
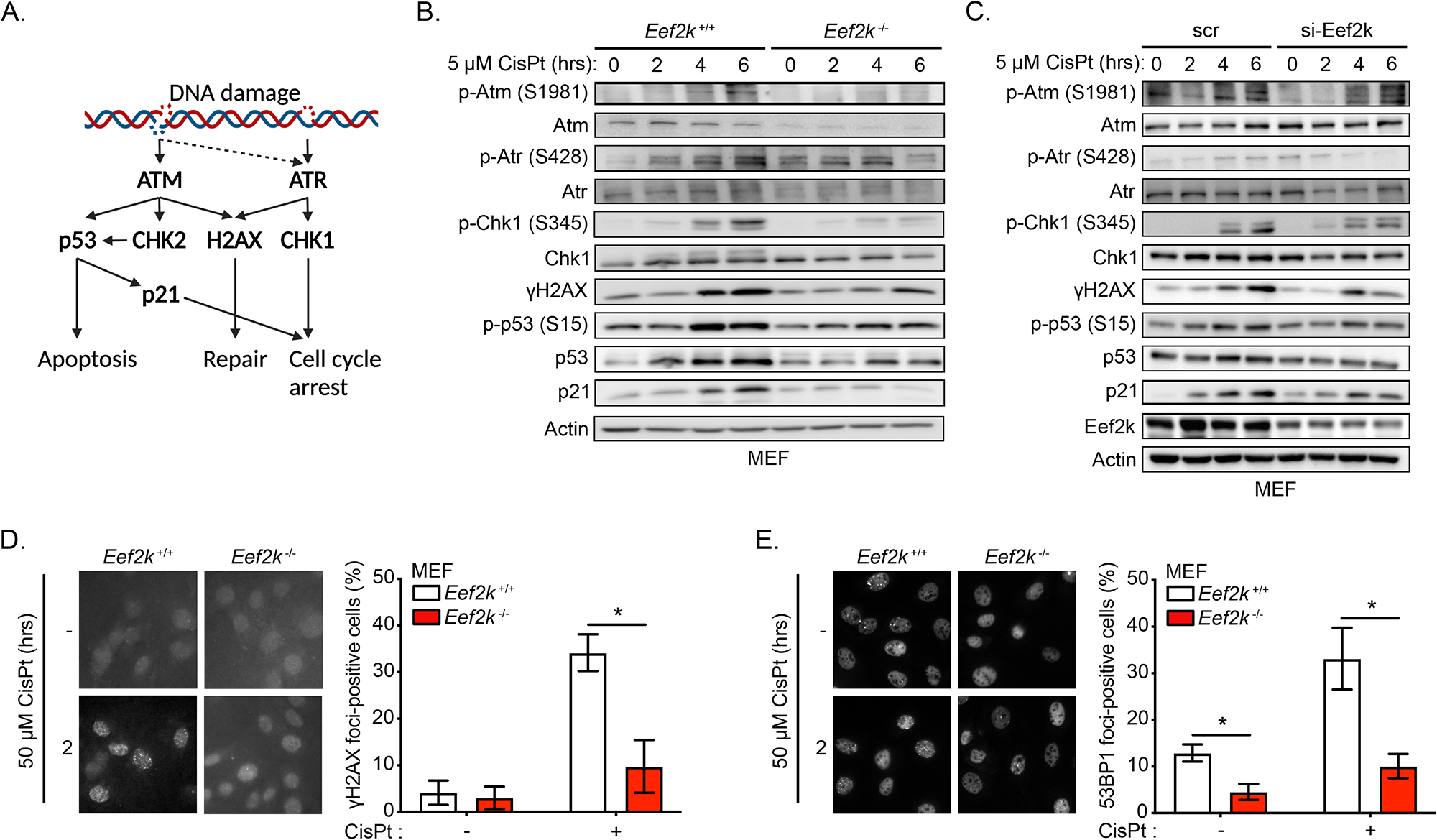
eEF2K supports the induction of the DDR pathways in response to cisplatin. **A.** Scheme illustrating ATM/ATR signalling in response to DNA damage. **B.** Level of activity of the ATM/ATR-dependent DDR pathways in *Eef2k*^+/+^ and *Eef2k*^-/-^ MEFs treated with CisPt (5 μM) for the indicated times, as measured with immunoblot analysis using the indicated antibodies. **C.** Level of activity of the ATM/ATR-dependent DDR pathways in MEFs transfected with siRNA targeting eEF2K (si-Eef2k) or scrambled control (scr) treated with CisPt (5 μM) for the indicated times, as measured with immunoblot analysis using the indicated antibodies. **D.** Level of γH2AX foci formation in *Eef2k*^+/+^ and *Eef2k*^-/-^ MEFs treated with CisPt (50 μM) or vehicle control for 2 hours, as determined by immunofluorescence using a γH2AX antibody (n = 3). **E.** Level of 53BP1 foci formation in *Eef2k*^+/+^ and *Eef2k*^-/-^ MEFs treated with CisPt (50 μM) or vehicle control for 2 hours, as determined by immunofluorescence using a 53BP1 antibody (n = 3). Data are expressed as mean ± SD; \**P* < 0.05.

The phosphorylation of H2AX as a downstream effector of ATM (and less commonly of ATR), leads to its early recruitment to DNA DSBs, while 53BP1 localization to these sites occurs later and is known to promote non-homologous end joining (NHEJ) repair by counteracting resection and therefore HRR. To further corroborate the above findings, we next examined the localization of γH2AX and 53BP1 to sites of DNA damage in cisplatin-treated *Eef2k*^+/+^ and *Eef2k*^-/-^ cells. Consistent with our previous data, while there was a significant increase in the number of *Eef2k*^+/+^ cells displaying γH2AX and 53BP1 foci following cisplatin treatment, *Eef2k*^-/-^ cells exhibited a clear deficiency in accumulation of these foci (Figures 2D-2E). Together, these data suggest that eEF2K promotes the induction of DNA damage-associated signalling in response to cisplatin.

### eEF2K promotes the efficient repair of DNA damage induced by cisplatin

Since *Eef2k*^+/+^ cells showed more proficient recruitment of γH2AX and 53BP1 to DNA damage lesions following cisplatin, as compared to *Eef2k*^-/-^ cells, we also wondered if these cells were better able to repair DNA damage following exposure to cisplatin. Thus, to measure the capacity of *Eef2k*^+/+^ and *Eef2k*^-/-^ cells to repair cisplatin-induced ICLs, we used a modified alkaline comet assay in which cells are acutely treated with cisplatin and allowed to recover for 2 or 24 hours. Following this recovery period, cells are subjected to X-ray irradiation immediately prior to harvesting, such that DNA with few cisplatin-induced ICLs will be fragmented by irradiation, migrating more rapidly and exhibiting a longer tail moment, while DNA fragments with multiple ICLs will remain connected through ICLs and therefore will migrate more slowly [16]. Control untreated cells displayed noticeably longer tail moments with no difference observed between *Eef2k*^+/+^ and *Eef2k*^-/-^ cells, which is indicative of a lack of crosslinked DNA, while cisplatin-treated cells that were allowed to recover for 2 hours displayed markedly diminished tail moments regardless of *Eef2k* status, consistent with highly crosslinked DNA in both *Eef2k*^+/+^ and *Eef2k*^-/-^ cell types (Figure 3A). Notably, following 24 hours of recovery post-cisplatin, *Eef2k*^-/-^ cells showed a significant decrease in tail moments relative to *Eef2k*^+/+^ cells, suggesting that these cells repaired cisplatin-induced ICL less efficiently. Similar differences were observed when sh-eEF2K cells were compared to sh-scr control cells (Figure 3B). Consistent with this, we also observed that *Eef2k*^+/+^ cells treated with cisplatin for 2 hours followed by a period of recovery showed a peak in the number of γH2AX foci at 6 hours, which was followed by a rapid decrease by 16 hours indicative of efficient DNA damage repair (Figure 3C; quantification in the right panel). In contrast, the number of γH2AX foci in *Eef2k*^-/-^ cells gradually increased and peaked at 16 hours and was only diminished at 24 hours, further suggesting that *Eef2k*^+/+^ cells were more efficient at repairing cisplatin-induced damage, particularly given the fact that γH2AX foci form during the excision step of ICL repair.

**Figure 3.**
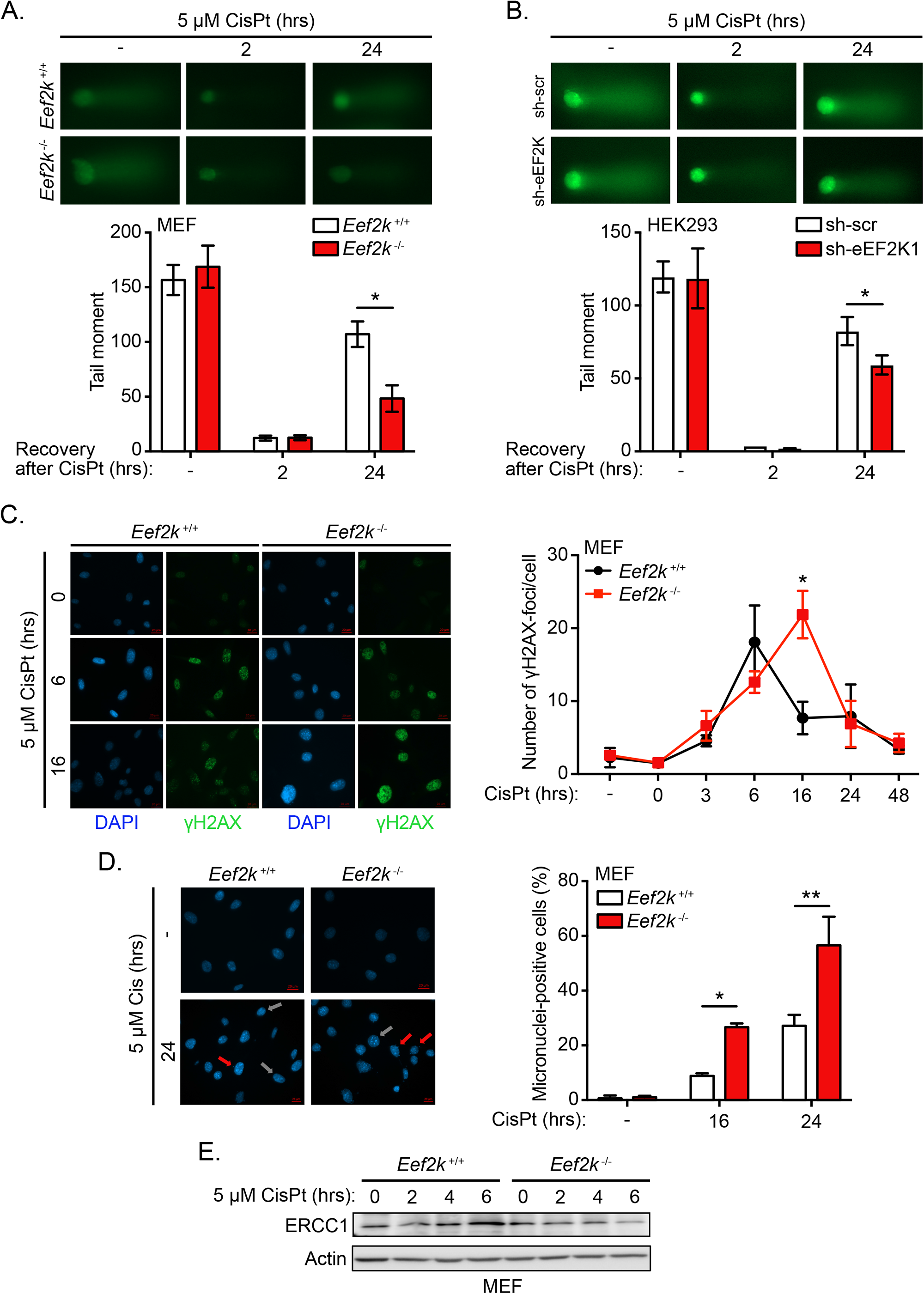
eEF2K promotes enhanced DNA damage repair in response to cisplatin. **A.** Level of DNA damage in *Eef2k*^+/+^ and *Eef2k*^-/-^ MEFs treated with vehicle (-) or with CisPt (5 μM) for 2 hours followed by 2 or 24 hours period of recovery, as measured by modified alkali comet assay (n = 3). **B.** Level of DNA damage in HEK293 cells stably expressing sh-eEF2K1 or sh-scr treated with vehicle (-) or with CisPt (5 μM) for 2 hours followed by 2 or 24 hours period of recovery, as measured by modified alkali comet assay (n = 3). **C.** Level of γH2AX foci formation and resolution in *Eef2k*^+/+^ and *Eef2k*^-/-^ MEFs treated with vehicle (-) or with CisPt (5 μM) for 2 hours followed by 0 to 48 hours period of recovery, as determined by immunofluorescence using a γH2AX antibody (n = 3). **D.** Micronuclei formation in *Eef2k*^+/+^ and *Eef2k*^-/-^ MEFs treated with vehicle (-) or with CisPt (5 μM) for 2 hours followed by 16 or 24 hours period of recovery, as measured using DAPI staining. Red arrows indicating micronuclei positive cells and grey arrows indicating micronuclei negative cells (n = 3). **E.** Level of ERCC1 expression in *Eef2k*^+/+^ and *Eef2k*^-/-^ MEFs treated with CisPt (5 μM) for the indicated times, as measured with immunoblot analysis using the indicated antibodies. Data are expressed as mean ± SD; \**P* < 0.05, \*\**P* < 0.01.

Deficient DNA repair can compromise chromatid separation during metaphase and lead to the accumulation of chromosomal abnormalities, such as aneuploidy, chromosomal breakage, and the formation of micronuclei, which indicate fragments of chromosomal DNA in daughter cells. Thus, we wondered whether deficient DNA repair in *Eef2k*^-/-^ cells leads to the accumulation of chromosomal abnormalities following cisplatin treatment. Indeed, we observed that at 16 and 24 hours post-cisplatin treatment, *Eef2k*^-/-^ cells displayed significantly higher numbers of micronuclei-positive cells (Figure 3D). Consistent with these findings, we also observed more *Eef2k*^-/-^ cells in S phase following cisplatin treatment as compared to *Eef2k*^+/+^ cells (Figure S2A). This may suggest either premature entry into S phase in spite of deficient ATM/ATR signalling, or may possibly be attributed to a slower progression of *Eef2k*^-/-^ cells through S phase due to stalled replication fork formation.

We next assessed the expression of ERCC1, a critical protein involved in ICL repair [13, 14, 15]. While *Eef2k*^+/+^ cells showed progressive accumulation of ERCC1 under cisplatin treatment for up to 6 hours, in contrast, we found that *Eef2k*^-/-^ cells failed to accumulate this protein under the same conditions (Figure 3E). Similar findings were observed in si-eEF2K and sh-eEF2K cells compared to scrambled controls (Figures S2B-S2C). Taken together, our findings suggest that eEF2K supports the induction of the DDR pathway, and enables more efficient DNA repair in response to cisplatin.

### The eEF2K-mediated cellular susceptibility towards cisplatin is p53-dependent

Our data provide evidence that while eEF2K enhances DNA damage repair under cisplatin treatment, paradoxically, our data also showed that eEF2K mediates increased susceptibility to cisplatin. To reconcile these observations, we hypothesized that while eEF2K deficient cells are less capable of mounting a DDR, they might in addition be defective in activating a p53-dependent pro-apoptotic program, resulting in lower susceptibility to cisplatin treatment. In support of this hypothesis, *Eef2k*^-/-^ cells displayed lower levels of active p53 (phosphorylation at Ser15), total p53, and p21 under a time course of 5 μM cisplatin (Figure 2B). Furthermore, compared to *Eef2k*^+/+^ cells, *Eef2k*^-/-^ cells also showed reduced active and total p53 over the same time course at higher cisplatin concentrations (50 μM), expected to induce greater levels of DNA damage (Figure 4A). Similar results were observed for HEK293 cells with stable eEF2K knockdown, which under 50 μM cisplatin treatment again showed significantly decreased accumulation of active and total p53, as well as noticeably decreased p21 levels, relative to control cells (scr) (Figure 4B). Of note, we observed that *Eef2k*^-/-^ cells had lower levels of *TP53* mRNA compared to *Eef2k*^+/+^ cells, either under basal conditions or 50 μM cisplatin treatment, suggesting that eEF2K can regulate p53 expression at the transcriptional level (Figure S3A), although the mechanism remains to be determined. To further corroborate this model, we subjected cells to cisplatin treatment in conjunction with siRNA-mediated knockdown of p53. Notably, knockdown of p53 rescued *Eef2k*^+/+^ cells from cisplatin-induced declines in cell viability across a range of cisplatin concentrations (Figure 4C and Figure S3B), but had no consistent effect on the viability of *Eef2k*^-/-^ cells under the same conditions (Figures 4D). Together, these findings suggest that while eEF2K plays a role in supporting an efficient DDR, it also facilitates p53-mediated cell death, likely in conditions when the damage surpasses a threshold of cellular DNA repair capacity.

**Figure 4.**
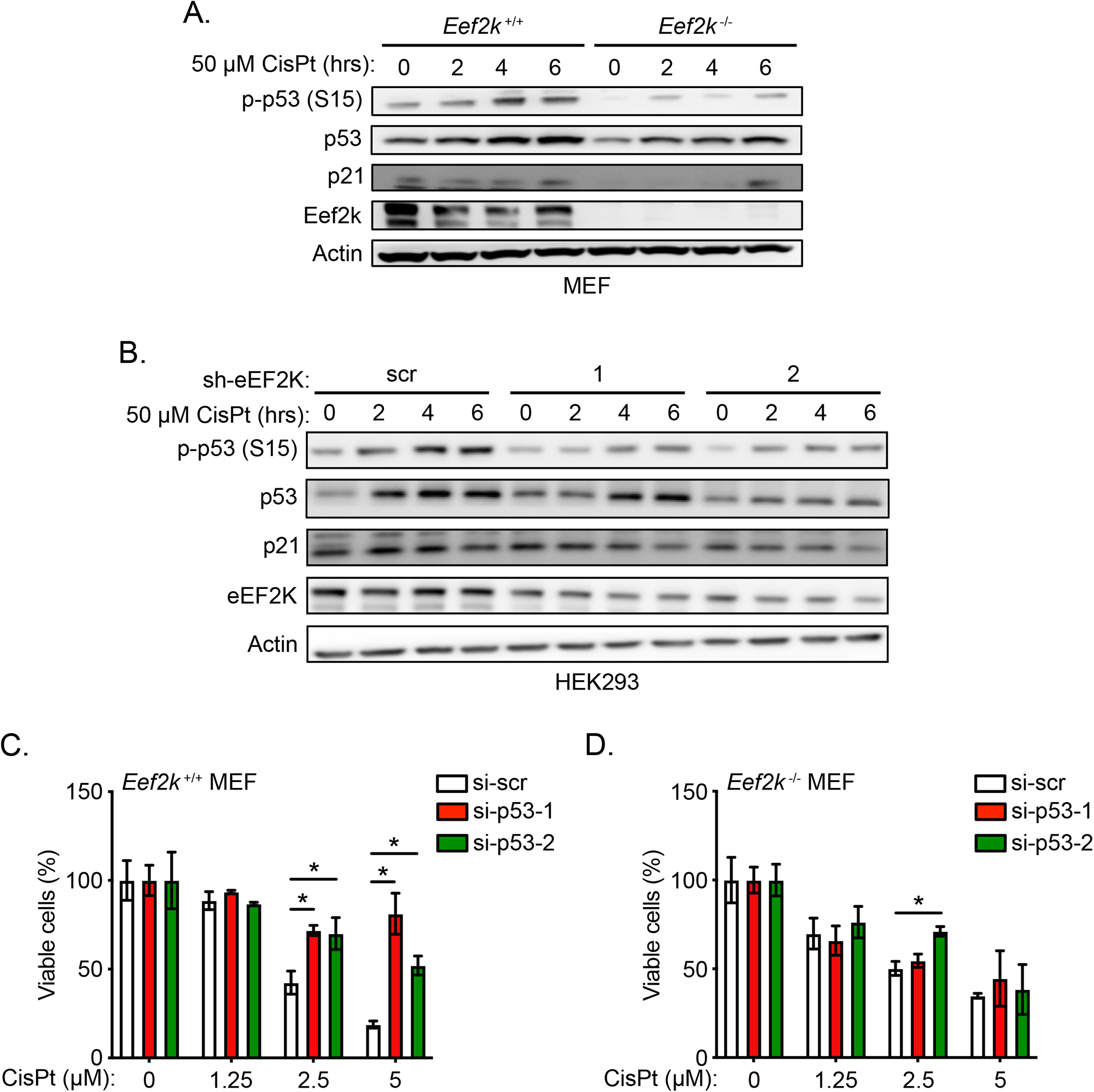
eEF2K-mediated cellular susceptibility towards cisplatin is dependent on p53 activation and expression. **A.** Level of p53 activation in *Eef2k*^+/+^ and *Eef2k*^-/-^ MEFs treated with CisPt (50 μM) for the indicated times, as measured with immunoblot analysis using the indicated antibodies. **B.** Level of p53 activation in HEK293 cells stably expressing sh-eEF2K1, sh-eEF2K2, or sh-scr treated with CisPt (50 μM) for the indicated times, as measured with immunoblot using the indicated antibodies. **C, D.** Viability of *Eef2k*^+/+^ MEFs (C) or *Eef2k^-/-^* MEFs (D) transfected with individual siRNA targeting p53 (si-p53-1 and si-p53-2) or scrambled control (si-scr) treated with the indicated concentrations of cisplatin (CisPt) or vehicle control for 48 hours, using an MTT assay (n = 3). Data are expressed as mean ± SD; \**P* < 0.05.

### The eEF2K ortholog, *efk-1*, promotes germ cell death in *C. elegans* in response to cisplatin

To ascertain if the role of eEF2K in DDR is recapitulated in vivo and conserved across species, we examined physiological responses to cisplatin using *C. elegans*. To this end, we used apoptosis and fertility assays to assess the response of wild type (*wt*) and *efk-1* (the *C. elegans* ortholog of *eEF2K)* mutant (*efk-1*) worms to cisplatin exposure. From this, we observed that *wt* worms had a higher number of corpses per gonad arm (Figure 5A), indicative of a higher rate of apoptosis, as well as a reduced number of hatched eggs per worm per hour (Figure 5B) relative to *efk-1* worms, at higher concentrations of cisplatin treatment. Interestingly, both *wt* and *efk-1* worms had no discernible differences in the proportions of viable eggs (Figure 5C). This suggests that while *wt* worms may respond appropriately to environmental stressors like DNA damage via the induction of apoptosis in germ cells or reducing the number of eggs laid, *efk-1* worms fail to mediate such a response. Thus, our data provide strong evidence that the eEF2K ortholog *efk-1* is a component of the DDR in *C. elegans*, arguing that this response is highly conserved.

**Figure 5.**
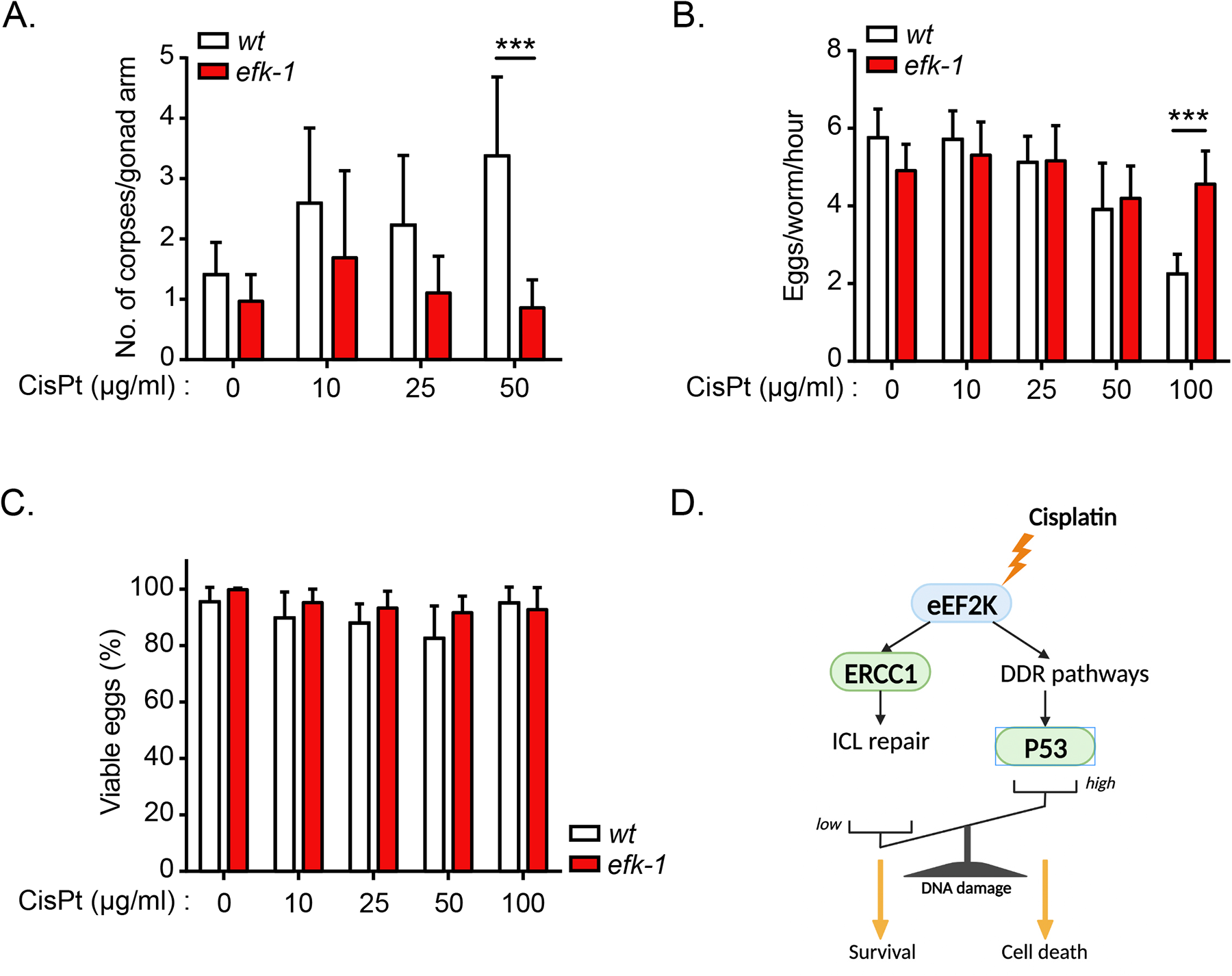
The eEF2K ortholog, *efk-1*, mediates germ cell death in *Caenorhabditis elegans* in response to cisplatin. **A.** Level of germ cell death in wild type (*wt*) and *efk-1(ok3609)* (*efk-1*) adult *C. elegans* treated with the indicated concentrations of CisPt for 48 hours, as measured by number of germ cell corpses per gonad arm. **B.** Egg production in wild type (*wt*) and *efk-1* knockout (*efk-1*) adult *C. elegans* treated with the indicated concentrations of CisPt for 48 hours, as measured by number of eggs produced per worm per hour. **C.** Percentage of viable eggs produced by *wt* and *efk-1* knockout (*efk-1*) adult *C. elegans* treated as in (B). Data are expressed as mean ± SD; \*\*\**P* < 0.001.

## DISCUSSION

DNA damage caused by cisplatin, in particular ICLs, triggers a coordinated DDR which results in DNA repair or, if the amount of DNA damage is excessive, leads to apoptosis induction. Here, we uncovered that eEF2K represents an unanticipated regulator of the DDR to cisplatin, acting through two distinct but potentially interconnected mechanisms. On the one hand, our data supports a model whereby eEF2K facilitates ICL repair by promoting ERCC1 expression, while on the other hand it promotes activation of DDR pathways, in turn inducing p53 expression/activity and leading to apoptosis as well as influencing the repair kinetics of ICL-induced damage. While such a dual function of eEF2K may seem paradoxical, the cellular output – cell death or cell survival – may directly depend on a balance between the amount of cisplatin-induced DNA damage and the intrinsic ability of cells to mount a pro-apoptotic p53 response (Figure 5D). It seems likely that eEF2K-deficient cells can tolerate a higher load of cisplatin-induced DNA damage without initiating apoptosis. In such a scenario, it could be expected that despite resisting cisplatin-induced cell death, eEF2K-deficient cells may accumulate more mutations and suffer increasing genomic instability in the long term, due to both the accumulation of DNA lesions and deficiency in coordinating error-free HRR, as expected from blunted ATM activation.

Previous reports have described a function for eEF2K in the DDR triggered by other DNA damage inducers, such as H_2_O_2_, doxorubicin and ionizing radiation [7, 8, 9]. In accordance with such findings, we characterized that in response to cisplatin, eEF2K is also activated, which in turn could restrict mRNA translation elongation as an integral part of the DDR, as reported for the response to doxorubicin [7]. In addition, our data highlight that eEF2K mediates apoptosis in response to cisplatin, further confirming earlier observations made with H_2_O_2,_ doxorubicin and ionizing radiation [8, 9], thus supporting the notion that eEF2K represents an important DDR mediator of apoptosis. While the mechanism underlying eEF2K’s pro-apoptotic function in response to H_2_O_2_ and doxorubicin was proposed to entail translational inhibition of transcripts encoding anti-apoptotic proteins, such as XIAP, c-FLIP and Mcl-1 [8], we demonstrated that eEF2K-mediated activation of p53 represents another mechanism.

Our data also suggests that eEF2K supports ICL repair, as eEF2K deficient cells harbor higher amounts of DNA damage following acute cisplatin treatment. In conjunction with the defective induction of the Chk-1 and p53 checkpoints in *Eef2k*^-/-^ cells, this could consequently lead to the observed accumulation of micronuclei in these cells. This is consistent with our findings that in response to cisplatin, *Eef2k*^-/-^ cells enter S phase and mitosis prematurely, albeit with compromised ATM and ATR signalling, enabling *Eef2k*^-/-^ cells to escape cell death and continue proliferating. We propose that in response to cisplatin, eEF2K coordinates cell cycle checkpoints and enhances ERCC1 expression, albeit through unknown mechanisms, together allowing cells to repair ICL (Figure 5D).

Our work reveals, unexpectedly, that eEF2K is required for activating the ATM and ATR signaling pathways in response to cisplatin. It is intriguing that eEF2K deficiency blunts the induction of the ATM and ATR pathways without preventing the build-up of DNA damage upon cisplatin treatment. Loss of eEF2K will release eEF2 from eEF2K-mediated inhibition, thus likely sustaining mRNA translation elongation activity in response to cisplatin, as was characterized for other stresses [3, 7]. Given that protein synthesis is the most ATP consuming process in the cell [17], failure to block mRNA translation elongation is expected to lead to ATP depletion in eEF2K deficient cells exposed to cisplatin, as shown in colon cancer cells under basal conditions [18]. However, this does not explain the lack of ATM activation in these cells as, on the contrary, ATM was reported to be induced by ATP depletion [19]. Another possibility is that loss of eEF2K may selectively promote the translation of transcripts encoding upstream inhibitors of ATM and/or ATR, including Repo-Man and WIP1 phosphatases [20, 21], which warrants further investigation.

We also uncovered that eEF2K promotes p53 expression and activity in response to cisplatin, highlighting eEF2K as a novel regulator of p53. While the mechanism involved is still elusive, apart from eEF2K-mediated activation of the upstream p53 regulators ATM and ATR, this regulation may explain the observation that *Eef2k*^-/-^ and p53^-/-^ cells display similar phenotypes when treated with cisplatin, ionizing radiation or doxorubicin, namely that eEF2K and p53 loss each confers resistance to these treatments [8, 9, 22, 23, 24]. Furthermore, such eEF2K regulation of p53 has broader physiological implications as it may provide an alternative mechanism for the ability of eEF2K to mediate apoptosis in oocytes, which contributes to preserve oocytes quality as reported [8]. Such a biological function of eEF2K was found to be conserved in the evolution, as characterized in *C. elegans* [8], in agreement with our findings that the *C. elegans* eEF2K ortholog *efk-1* mediates apoptosis in response to cisplatin similarly to mammalian eEF2K. However, in contrast to Chu *et al* [8], we did not observe a difference in the apoptosis rates, as measured by number of corpses, between wild type and *efk-1(ok3609)* worms under basal conditions but only upon high concentrations of cisplatin. It is worth noting that in response to ionizing radiation it was reported that eEF2K mediates cell death even in p53 deficient cells [9], suggesting that how eEF2K function in the DDR is both p53-dependent and independent, which may depend on the type of DNA damage inducers and the extent of damage. Since eEF2K expression and activity is deregulated in cancers, eEF2K regulation of p53 may have clinical implications in cancer [3, 4]. Indeed, by supporting p53 activation in p53 functional cancers, eEF2K may sensitize tumor cells to cisplatin, which is commonly used as a chemotherapy agent in various tumor types. With that perspective, levels of eEF2K expression and activity could serve as biomarkers for tumors that would best respond to cisplatin treatment. However, the role of eEF2K in cancer is still being scrutinized, as eEF2K displays both pro-tumorigenic and tumor suppressive functions depending on the tumor entities and the type of stress [25]. Since activation of the DDR acts as a barrier against tumorigenesis [26, 27, 28], eEF2K activation of DDR pathways could support a tumor suppressive function of eEF2K. However, the DDR can also act as an Achilles heel in some cancers; a large number of novel mono- and combination therapies utilising inhibitors against cell cycle and DDR enzymes are currently undergoing clinical trials [29]. Thus, by facilitating ICL repair, eEF2K may allow tumor cells to tolerate cisplatin treatment at certain doses, which could in the long term lead to the emergence of cisplatin-resistant tumor clones. This also warrants further investigation.

Overall, our study reveals that the mRNA translation regulator eEF2K represents a novel evolutionary conserved regulator of the DDR to cisplatin, which controls the rates of apoptosis and of DNA repair, depending on the extent of DNA damage, by regulating the ATM-ATR-p53 pathways and ERCC1 levels, respectively.

## MATERIALS AND METHODS

### Cell culture and cisplatin treatment

Mouse embryonic fibroblasts (MEFs) and human embryonic kidney 293 cells (HEK293) were obtained from ATCC and maintained in Dulbecco’s modified Eagle medium (DMEM; Invitrogen) supplemented with 10% fetal bovine serum (FBS). All cell lines were maintained at 37°C in an atmosphere of 5% CO_2_. The immortalized *Eef2k*^+/+^ and *Eef2k*^-/-^ MEFs were gifts of Dr. Alexey Ryazanov (University of Medicine and Dentistry of New Jersey). Cisplatin (15663-27-1, Sigma-Aldrich) was resuspended in DMSO at 50 mM concentration. The cisplatin aliquots were stored at -80°C for a maximum period of 3 months. Each aliquot was used only once after thawing.

### *C. elegans* cultivation and cisplatin treatment

*C. elegans* strains N2 (wild type) and *efk-1* (*ok3609*) deletion mutant (RB2588) were obtained from the Caenorhabditis Genetics Center funded by the NIH Office of Research Infrastructure Programs (P40 OD010440) and cultivated at 20°C on Nematode Growth Medium (NGM) plates seeded with Escherichia coli OP50, as described in [30]. Cisplatin-containing NGM plates were prepared by adding cisplatin (Sigma Aldrich) dissolved in 150 mM NaCl to autoclaved and cooled NGM at 55°C. The medium was then poured into petri dishes and following overnight solidification,*E. coli* OP50 were added and allowed to grow for an additional 2 days at room temperature. After synchronization of the animals by egg prep (1% NaClO and 0.4 M NaOH for 5 minutes), semi-synchronous eggs were placed on NGM plates without cisplatin and grown for 3 days at 20°C to reach adulthood. From the third day after synchronization, the animals were treated with cisplatin using cisplatin-containing NGM plates.

### *C. elegans* germline apoptosis assay and reproduction/embryonic viability assays

Acridine Orange was used for staining to detect apoptotic cells in the germline, as described in [30]. After anesthetizing the nematodes with 15 mM sodium azide, stained germline corpses were examined using an Olympus CJX41 microscope in the GFP channel, using 400X magnification. The quantification of stained apoptotic cells was conducted in the posterior gonad arm. To measure reproduction and embryonic viability, 3-day old worms were treated with cisplatin for 24 hours. Next, 3 worms per experimental group were transferred to small NGM plates that did not contain cisplatin and were incubated for 3-4 hours. Following this, the adult worms were removed and the number of eggs was counted (reproduction). After an additional day, the number of hatched larvae and remaining eggs on the plates were recorded to determine the ratio of hatched larvae to the total number of eggs laid (embryonic viability).

### Short hairpin RNA and lentivirus production

Lentivirus was produced by transfecting HEK293T cells grown in T25 flasks with 3 μg of pLK0.1 plasmid containing relevant shRNA sequences (Sigma-Aldrich) together with 1.5 μg and 3 μg of lentiviral envelope and packaging vectors pVSVG and psPAX2, respectively. Viral media was collected and filtered 24 hours, 48 hours, and 72 hours post transfection. 1:1 viral supernatant to fresh media was used to infect HEK293 cells. eEF2K expression was depleted in HEK293 cells using sheEF2K lentiviral pLK0.1 plasmids containing the targeting sequences (Sigma-Aldrich): sh-eEF2K-1-CCG GCC ACT CAT ACA GTA ATC GGA ACT CGA GTT CCG ATT ACT GTA TGA GTG GTT TTT and sh-eEF2K-2-CCG GCG ATGA GGA AGG TTA CTT CAT CTC GAG ATG AAG TAA CCT TCC TCA TCG TTT TT. As controls, HEK293 cells were transduced with pLK0.1 lentiviral plasmid containing scrambled non-targeting sequence (Sigma-Aldrich). Puromycin (5 µg/mL) was used for selection.

### siRNA Transfection

Cells were plated at ∼15% confluency and transfected with 25 nM siRNA using RNAi Lipofectamine (Invitrogen). Cells were incubated for 48 to 72 hours before being treated with cisplatin. Stealth RNAi negative control duplexes, medium GC Duplex (Invitrogen), was used as control for all the RNAi experiments. Pooled mouse eEF2K siRNA (sc-39011, Santa Cruz) was used to knockdown eEF2K expression. siRNAs-mediated knockdown P53 expression was performed using mouse p53 siRNAs with the following sequences si-p53-1 GUA AAC GCU UCG AGA UGU U and si-p53-2 AAA UUU GUA UCC CGA GUA U (Dharmacon).

### Immunofluorescence Staining

Cells were plated on coverslips one day prior to the treatment. At end point, the cells were fixed in 4% paraformaldehyde (PFA) in phosphate buffered saline (PBS) for 15 minutes. PFA was washed off and coverslips were stored in PBS at 4°C overnight. Cells were permeabilized by incubation in 0.2% Triton X-100 in PBS (T-PBS) for 30 minutes, followed by incubation in blocking solution 5% bovine serum albumin (BSA; Thermo-Fisher) in 0.2% T-PBS for 30 minutes. Cells were incubated with primary antibodies for 1 hour at room temperature. The primary antibodies were washed off 3 times 10 minutes with 0.2% T-PBS. Cells were incubated for 1 hour with the secondary antibody. Cells were washed 3 times 10 minutes with 0.2% T-PBS. Cover slips were mounted on slides with mounting media containing DAPI (Vectashield) and sealed with clear nail polish.

### Immunofluorescence microscopy

Zeiss Axio Colibri fluorescent microscope was used to acquire images. Foci of 150 cells were quantified for each replicate. ImageJ software was used to detect the maximal points of fluorescence per nucleus and the average number of foci per nucleus per biological replicate were measured. DAPI nuclei stain was used to visualize micronuclei. The data is represented as the mean value of the three independent replicates.

### Immunoblot analysis

Whole cell lysates were collected using the lysis buffer (10 mM HEPES pH 7.4, 50 mM KCl, 5 mM MgCl_2_, and 0.5% NP-40). Protein concentration was measured and lysates were boiled in 1X SDS loading buffer for 10 minutes. Samples were run on SDS-PAGE gel and transferred to nitrocellulose membranes. The membrane was blocked in 5% milk in TBS Tween 0.1% (TBST) for 30 minutes and incubated over night at 4°C with the appropriate primary antibody in 2.5% milk in TBST. Following 3 washes in TBST, the membranes were incubated for 1 hour at room temperature with the secondary antibody in 2.5% milk in TBST, followed by 3 washes in TBST. The Pierce™ ECL Plus Western Blotting Substrate (Pierce, Thermo Scientific) was used for signal detection.

Primary and secondary antibodies for immunoblotting and immunofluorescence were: 53BP1 (sc-515841, Santa Cruz), ATM (2873, Cell Signaling), p-ATMser1981 (05-740, Millipore), ATR (13934, Cell Signaling), p-ATRthr1989 (30632, Cell Signaling), Actin (sc-2354, Santa Cruz), Caspase 3 (9662, Cell Signaling), PARP (9542, Cell Signaling), Chk1 (2360, Cell Signaling), p-Chk1ser345 (2348, Cell Signaling), ERCC1 (OAAF01786, Aviva Systems Biology), p21 (sc-6246, Santa Cruz), p53 (2527, Cell Signaling), p-p53ser15 (9284, Cell Signaling), H2AX (7631, Cell Signaling), γH2AX (9718, Cell Signaling), eEF2K (ab46787, Abcam), eEF2 (2332, Cell Signaling), p-eEF2thr56 (2331, Cell Signaling), and Alexa Fluor™ 488 (11008, Invitrogen).

### Modified alkaline comet assay

A modified alkaline comet assay procedure was conducted as previously described [16]. Cells were treated with 5 μM cisplatin for 2 hours, followed by 2X PBS washes. Cells were incubated with fresh media for 0, 2 or 24 hours. Cells were treated with 5 Grays of X-ray radiation and collected immediately by trypsinization. About 15,000 cells were suspended in 1% agarose gel (4018, Sigma) and embedded on a microscope slide. Cells were lysed overnight at 4°C in alkaline lysis solution. Lysis solution was washed off and electrophoresis was carried out in detergent free alkaline solution for 20 minutes at 20 V. The samples were stained with PI solution. Comet images were captured using a Zeiss Axio Colibri fluorescent microscope. Each condition was carried out in triplicate, and 50-100 comets were measured per replicate. The software Casplab was used to measure the comet tail moment.

### Trypan blue cell death assay

Cells were plated at a confluency of 30% in 6-well or 12-well plates and treated with 0-10 μM cisplatin for 48 hours. All cells were collected, washed and resuspended in PBS. Cells were mixed with trypan blue 0.4% solution at a 1:1 concentration and 10 µL of the mixture was applied to the hemocytometer. The total number of cells and trypan blue positive dead cells were counted. The percentage of dead cells was measured as the number of trypan blue positive cells over the total number of cells.

### MTT Assay

Cells were plated at a confluency of 30% in 6 well or 12 well plates. MEFs and HEK293 cells were treated with 0-20 μM cisplatin for 48 hours or 72 hours, respectively. Media containing the drug was removed, and cells were incubated with MTT (0.25 mg/mL) in fresh media for 3-4 hours. Medium was removed and 1 mL DMSO was added to cells to dissolve the formazan crystals. After crystals had fully dissolved, 80 µL of the dissolved solution was added to a 96 well plate, and absorbance was measured by a SpectaMax i3 plate-reader (Molecular Devices) at an absorbance value of 590 nm and an absorbance value of 690 nm was used as a control.

### Quantitative RT-PCR

Wild type and eEF2K knockout MEFs were treated with 50 µM of cisplatin or DMSO for 2 to 6 hours. Cells were washed with PBS and total RNA was extracted using RNeasy kit (Qiagen). cDNA was synthesized using High-Capacity cDNA reverse transcription kit (Applied Biosystems). The cDNA, SYBR Green mix, and the primers were mixed according to manufactures instructions. The samples were run on the Quant Studio 6 Flex RT-PCR system. The following TP53 primer sequences were used to amplify the TP53 cDNA: forward 5’-ACG CTT CTC CGA AGA CTG G – 3, and reverse 5’-AGG GAG CTC GAG GCT GAT A -3’ and the actin cDNA was amplified by using the following primer sequences: forward 5’-GTG ACG TTG ACA TCC GTA AAG A-3’ and reverse 5’-GCC GGA CTC ATC GTA CTC C-3’ (IDT). The qRT-PCR results were analyzed comparing the cycle threshold (CT) values of TP53 to the CT values of the reference gene, actin.

### Flow cytometry cell cycle profile analysis

Cell cycle analysis procedure was conducted as previously described [31]. The cells were plated on 4 cm plates at a 60% confluency. Cells were treated with 5 μM cisplatin for 0-24 hours, and one hour prior to collection cells were treated with BrdU at a final concentration of 10 µM to label cells in S-phase. Cells were collected with trypsinization and re-suspended in 500 µL of PBS. Cells were fixed in 70% ethanol and stored at -20°C. Ethanol was washed off and cells were resuspended in PBS. BrdU-FITC conjugated antibody (BD Biosciences) was used to label the S-phase cells and PI to label DNA content. Flow cytometry analysis was carried out on BD FACSCalibur. FlowJo software was used to analyze the results.

### Quantification and Statistical Analysis

All experiments were, if not otherwise stated, independently carried out at least three times. Statistical significance was calculated using Student’s t-test in GraphPad Prism 8. The data are represented as means +/-standard deviation. A p-value of less than 0.05 was considered to be significant.

### Resource Availability

Further information and requests for reagents may be directed to, and will be fulfilled by the corresponding authors.

## ACKNOWLEDGEMENTS

This work was supported in part by funds from CIHR Foundation grant FDN-143280 (to P.H.S.). G.L. was supported by a grant from the German Cancer Aid (grant no. 70112624).

## DECLARATION OF INTERESTS

The authors declare no competing interests.

## SUPPLEMENTARY FIGURE LEGENDS

**Figure S1.**
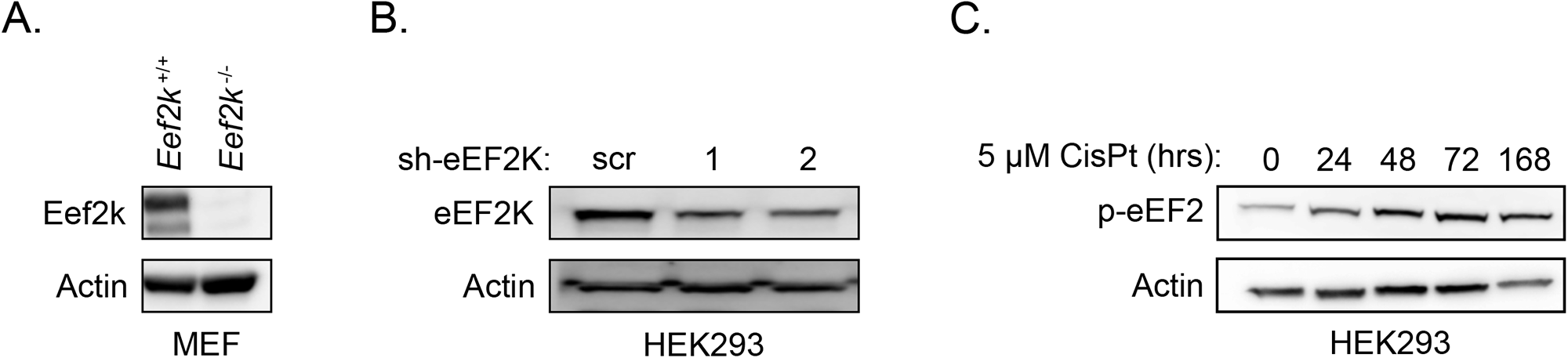
Validation of eEF2K knockout and knockdown, and levels of eEF2 phosphorylation under cisplatin treatment. **A.** eEF2K expression levels in *Eef2k*^+/+^ and *Eef2k*^-/-^ MEFs, as measured with immunoblot analysis using the indicated antibodies. **B.** eEF2K expression levels in HEK293 cells stably expressing individual shRNAs targeting eEF2K (sh-eEF2K1 and sh-eEF2K2) or scrambled control (sh-scr), as measured with immunoblot analysis using the indicated antibodies. **C.** Level of eEF2 phosphorylation (p-eEF2) in HEK293 cells treated with CisPt (5 μM) for the indicated times, as measured with immunoblot analysis using the indicated antibodies.

**Figure S2.**
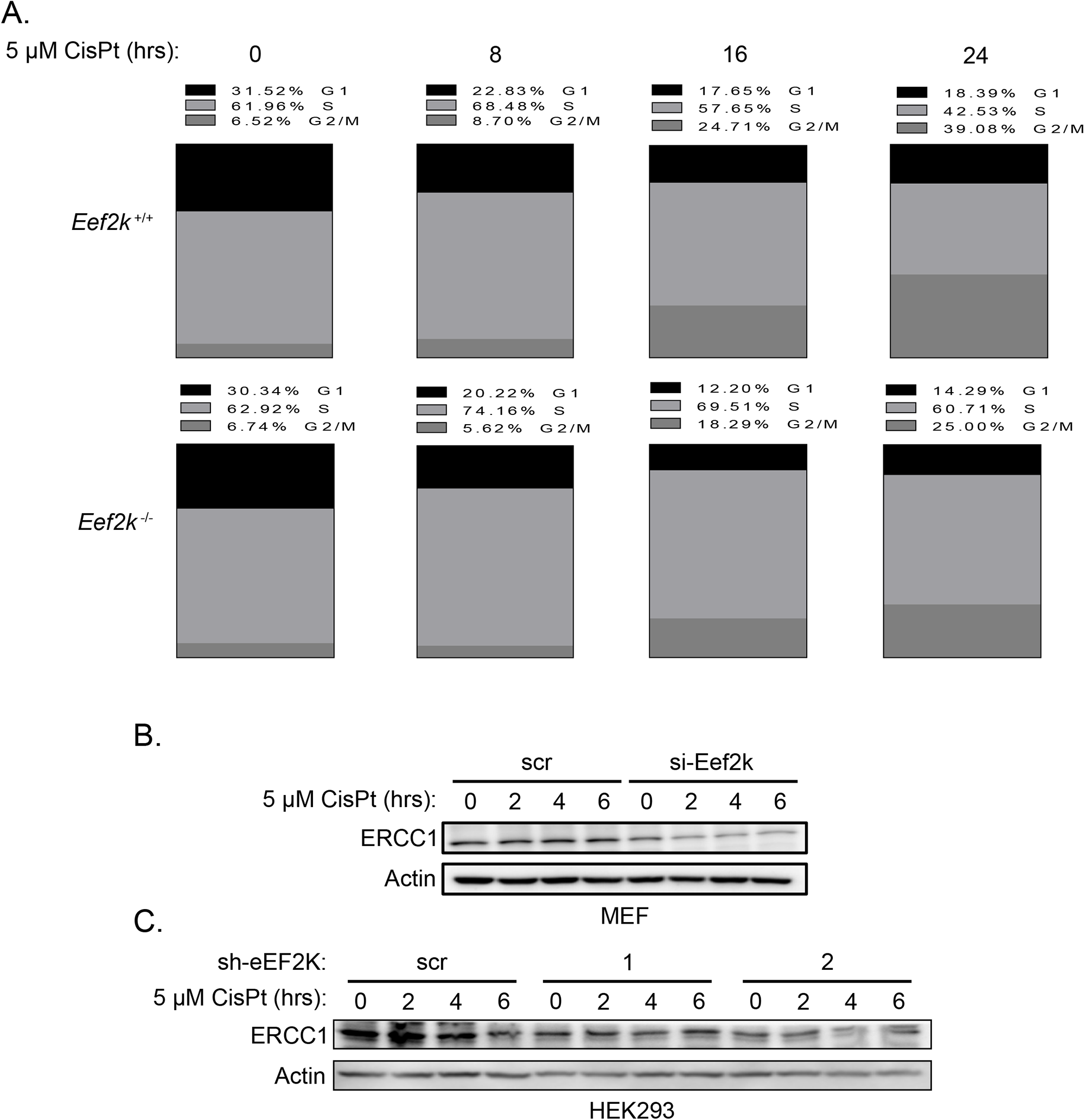
Cell cycle analysis and ERCC1 expression of eEF2K-deficient cells versus control cells under cisplatin treatment. **A.** Cell cycle analysis of *Eef2k*^+/+^ and *Eef2k*^-/-^ MEFs treated with CisPt (5 μM) for the indicated times, as measured by propidium iodide (PI) staining. **B.** ERCC1 expression levels in MEFs transfected with siRNA targeting eEF2K (si-Eef2k) or scrambled control (scr) treated with CisPt (5 μM) for the indicated times, as measured with immunoblot analysis using the indicated antibodies. **C.** ERCC1 expression levels in HEK293 cells stably expressing individual shRNAs targeting eEF2K (sh-eEF2K1 and sh-eEF2K2) or scrambled control (sh-scr) treated with CisPt (5 μM) for the indicated times, as measured with immunoblot analysis.

**Figure S3.**
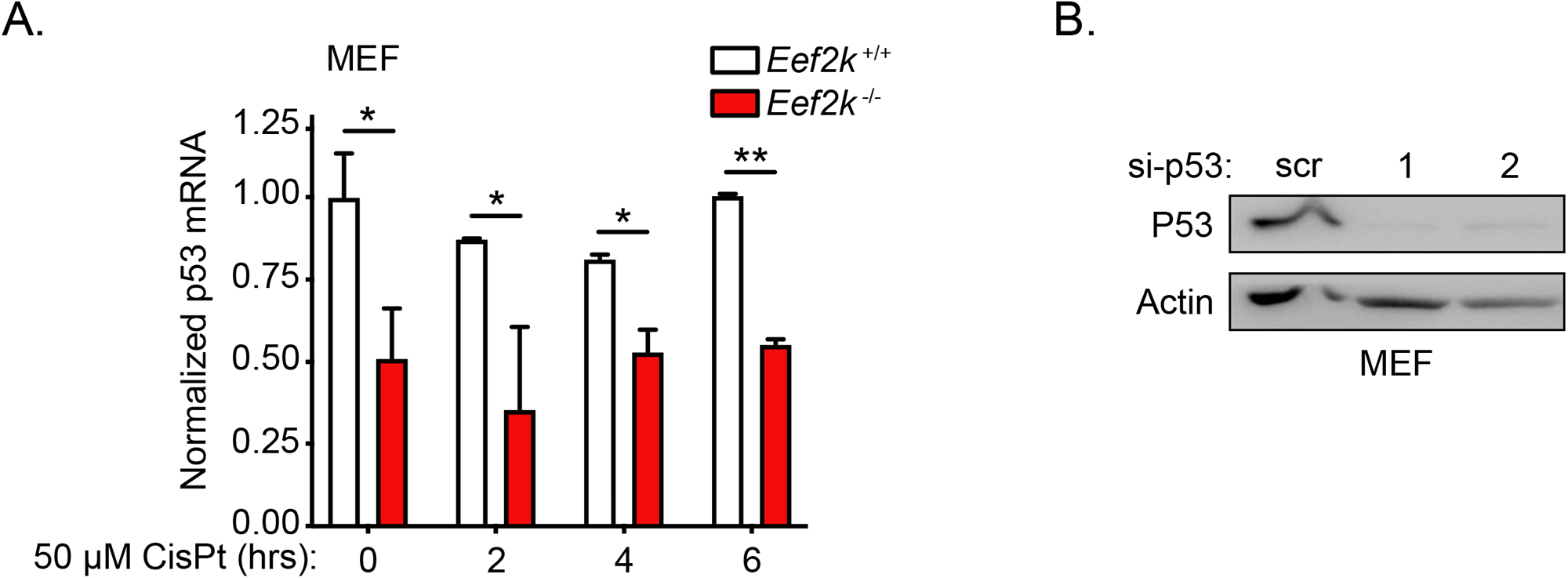
p53 mRNA levels of eEF2K-deficient and control MEFs under cisplatin treatment. **A.** Normalized p53 mRNA levels in *Eef2k*^+/+^ and *Eef2k*^-/-^ MEFs treated with 50 μM CisPt for the indicated times, as measured with qRT-PCR. **B.** Level of total p53 protein in *Eef2k*^+/+^ cells transfected with siRNA targeting p53 (si-p53-1 and si-p53-2) or scrambled control (scr), as measured with immunoblot analysis using the indicated antibodies. Data are expressed as mean ± SD; \**P* < 0.05, \*\**P* < 0.01.

## Notes

### Competing Interest Statement

The authors have declared no competing interest.

